# Spatiotemporal heterogeneity and the long-term impact of meteorological, environmental and socio-economic factors of scrub typhus in China from 2012 to 2018

**DOI:** 10.1101/2022.05.30.493950

**Authors:** Jiaojiao Qian, Changqiang Zhu, Heng Lv, Hongliang Chu, Ji He, Chongcai Wang, Yong Qi, Yizhe Luo, Na Yue, Yifan Wu, Fuqiang Ye, Jiying, Chunhui Wang, Weilong Tan

## Abstract

Large-scale outbreaks of scrub typhus combined with the emergence of this vector-borne rickettsiosis in new areas indicate that this disease remains seriously neglected. This study aimed to explore the long-term changes and regional leading factors of scrub typhus in China, so as to provide fresh insights for the prevention and control of this disease. In this study, a Bayesian space-time hierarchical model (BSTHM) was used to identify the long-term spatiotemporal heterogeneity of scrub typhus and quantify the association between meteorological factors and scrub typhus in southern and northern China from 2012 to 2018. GeoDetector model was used to quantify the dominant forces of environmental and socioeconomic factors in the Northern and the Southern China. Scrub typhus often appeared in summer and autumn (June to November), and epidemically peaked in October, with obvious temporal seasonality. Spatially, the hot spots (high-risk regions) were concentrated in the south, on the contrary the cold spots (low-risk regions) in the north. In addition, the main meteorological factor, average temperature, gave a significant impact in both areas. The average temperature increased by 1 °C, resulting in a decrease of 1.10% in southern China and an increase of 0.96% in northern China in the risk of scrub typhus. The determinant environmental and socio-economic factors of scrub typhus in the two areas were altitude and per capita GDP, with q-values of 0.91 and 0.87, respectively. Meteorological, environmental and socio-economic factors had a significant impact on the distribution of scrub typhus, with obvious seasonality and spatial heterogeneity. This study provides helpful suggestions and basis for reasonably allocating resources and controlling the occurrence of scrub typhus.

**Author summary:** Scrub typhus is a natural-focus disease caused by the bite of chigger mite larval. In this study, we use BSTHM to capture the overall temporal trend and spatial hot spots of scrub typhus, and quantify the relationship between the disease and major meteorological factors. Meanwhile, Geodetector model was used to quantify the influence of other potential risk factors and estimate the spatio-temporal heterogeneity of scrub typhus. The results showed that scrub typhus had significant seasonality, with a q value of 0.52, and spatial heterogeneity, with a q-value of 0.64. Scrub typhus mainly occurred in summer and autumn, and high-risk areas were mainly distributed in southern China (Yunnan, Hainan and Guangdong). These heterogeneity were closely related to the vector and host. Whether in the South or the north, scrub typhus was closely related to risk factors such as temperature, per capita GDP, NDVI, altitude and the percentage of children aged 0-14. These results suggest that the relevant departments should strengthen the monitoring of the ecological environment, the host and vector of Orientia tsutsugamushi, and strengthen the risk awareness, so as to prevent and control the possible increased risk of scrub typhus under these meteorological, environmental and socio-economic conditions. Considering the differences in different regions, resources should be allocated reasonably.

## 1. Introduction

Scrub typhus is a natural-focus disease caused by Orientia tsutsugamushi, which is mainly carried by rodents and chigger mites, and transmitted through the bite of an infected chigger mite larval occasionally. This disease is characterized of fever, eschar, rash and lymph node enlargement, and can even lead to multiple organ failure and death in severe cases with a fatality rate of up to 70%(Wongsantichon et al., 2020). O.tsutsugamushi is geographically endemic across vast areas of Asia and islands in the Pacific and Indian Oceans. Scrub typhus is one of the most widespread and severe infections of rickettsia which lead to more than one million infective cases each year(Tshokey et al., 2017; Wongsantichon et al., 2020). An increasing number of research(Izzard et al., 2010; Luce-Fedrow et al., 2018; Masakhwe et al., 2018; Weitzel et al., 2016; Weitzel et al., 2019) indicated that the distribution of scrub typhus was no longer limited to the Asia-Pacific region, but is spreading worldwide. Without an effective vaccine and rapid diagnosis, scrub typhus remains a severe public health threat.

In China, scrub typhus was always confined to tropical and subtropical areas of southern China for a long time period after it was identified in Yunnan province in 1943(Fan et al., 1987). The first case of autumn-winter tsutsugamushi occurred in Mengyin County, Shandong Province in 1986, indicating its geographical diffusion to the northern China (Liu et al., 2009). Since then, new natural foci have been reported and identified continuously, resulting in a significant increase in geographical distribution and a remarkable upward trend of the case number annually. Some studies indicated that spatiotemporal heterogeneity of scrub typhus was initially observed between the southern and northern China(Li et al., 2020; Yao et al., 2019).

Numerous studies(Brugueras et al., 2020; Mala and Jat, 2019; Miao et al., 2021; Parham et al., 2015; Zheng et al., 2019) have found that vector-borne diseases were not only related to environmental factors such as meteorological conditions, but also influenced by socio-economic factors. For example, Zheng et al.’ study (Zheng et al., 2019) revealed that in the southern China, relative humidity(RH), temperature, normalized vegetation index (NDVI) and altitude were the main factors affecting the spreading of scrub typhus. However, few studies have estimated the temporal and spatial heterogeneity of the scrub typhus risk and quantified the impact of environmental, socio-economic, and meteorological factors in southern and northern China from a long-term perspective. This study aimed to explore the temporal and spatial heterogeneity of the incidence of scrub typhus and reveal the cold and hot spots at provincial level from 2012 to 2018. We also sought to quantify the impact of environmental, socio-economic, and meteorological factors so as to provide novel insights into the health threat posed by scrub typhus.

## 2. Methods

### 2.1 Ethics statement

This study was approved by the ethics committee of Nanjing Bioengineering (Gene) Technology Center for Medicines(No:2021BY07).Patient consent was not required because no patients’ individual information was included in this study and population data were collected from the public database of China.

### 2.2 Study region

The geological environment, agricultural production, natural conditions, living customs, and socio-economic conditions are obviously different on the south and north regions divided by the Qinling-Huai River Mountains in China(Fig.1). The differences between the north and the south, such as climate and geological environment, et al, result in the significant difference in the species and distribution of vectors, host animals and the pathogens. Therefore, in our study, China was divided into two study areas: southern (Tibet, Jiangsu, Anhui, Hubei, Zhejiang, Jiangxi, Hunan, Yunnan, Guizhou, Fujian, Guangxi, Guangdong, Hainan, Chongqing, Shanghai, and Sichuan) and northern (Beijing, Hebei, Ningxia, Henan, Shandong, Inner Mongolia, Shanxi, Shaanxi, Gansu, Qinghai, Tianjin, Liaoning, Jilin, Xinjiang, and Heilongjiang).

**Fig. 1.**
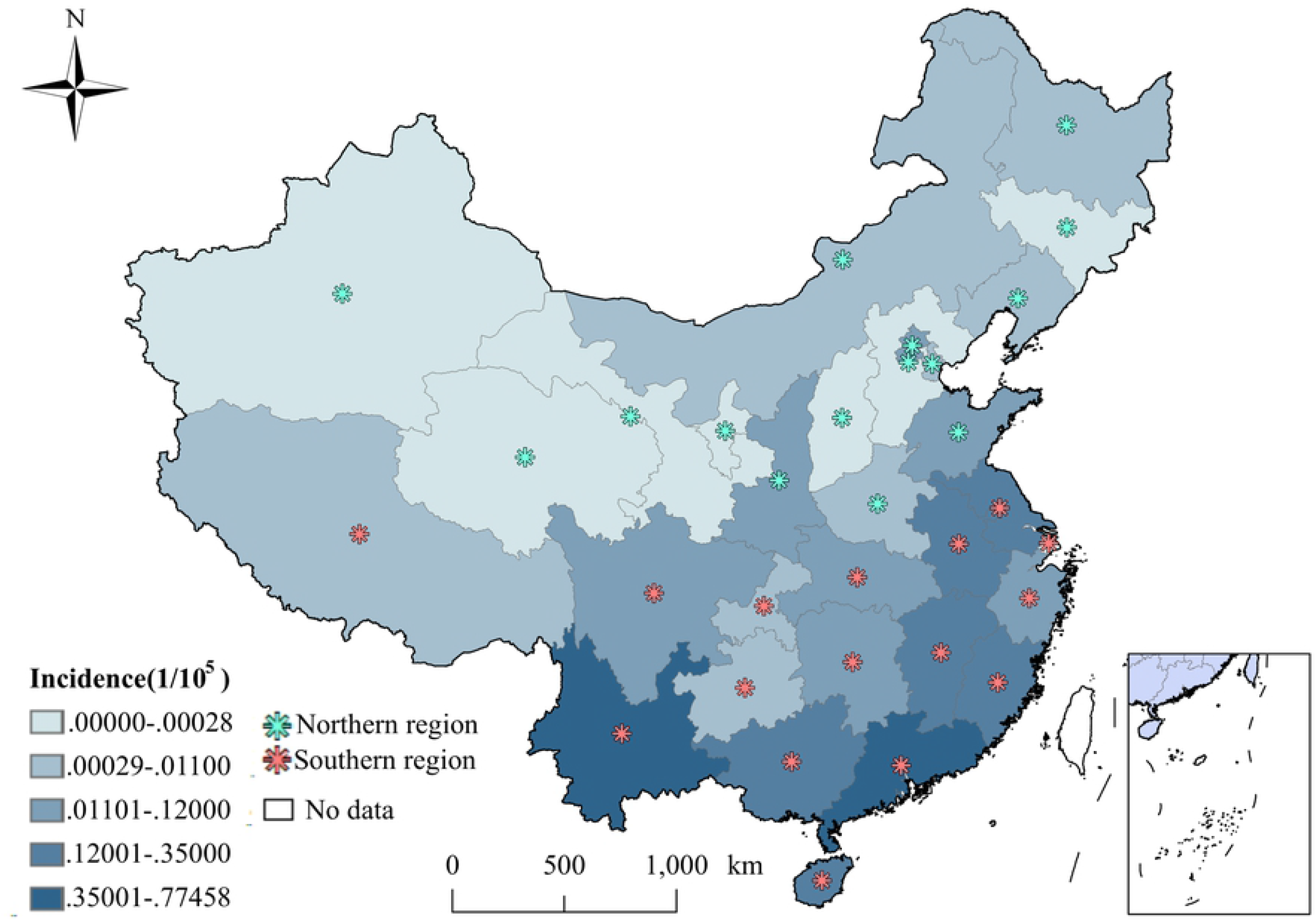
Monthly incidence of scrub typhus between 2012 and 2018 in China.

### 2.3 Data source

Monthly data of scrub typhus incidence for the period from January 2012 to December 2018 were obtained from the Chinese Center for Disease Control and Prevention (https://www.phsciencedata.cn/Share/). Monthly meteorological data, including average temperature, precipitation, RH, and hours of sunshine, and yearly province-level socioeconomic variables data, including per capita GDP, population density, number of health technicians per 1000 persons, illiteracy rate, number of medical beds per 1000 persons, percentage of population aged 0-14, and percentage of population over 65, were acquired from the Chinese economic Statistical Yearbook(http://www.stats.gov.cn/). The average data of provincial-level altitude and normalized Difference Vegetation Index (NDVI) were collected from the Geospatial data cloud(http://www.gscloud.cn/)(Fig.2).

**Fig. 2.**
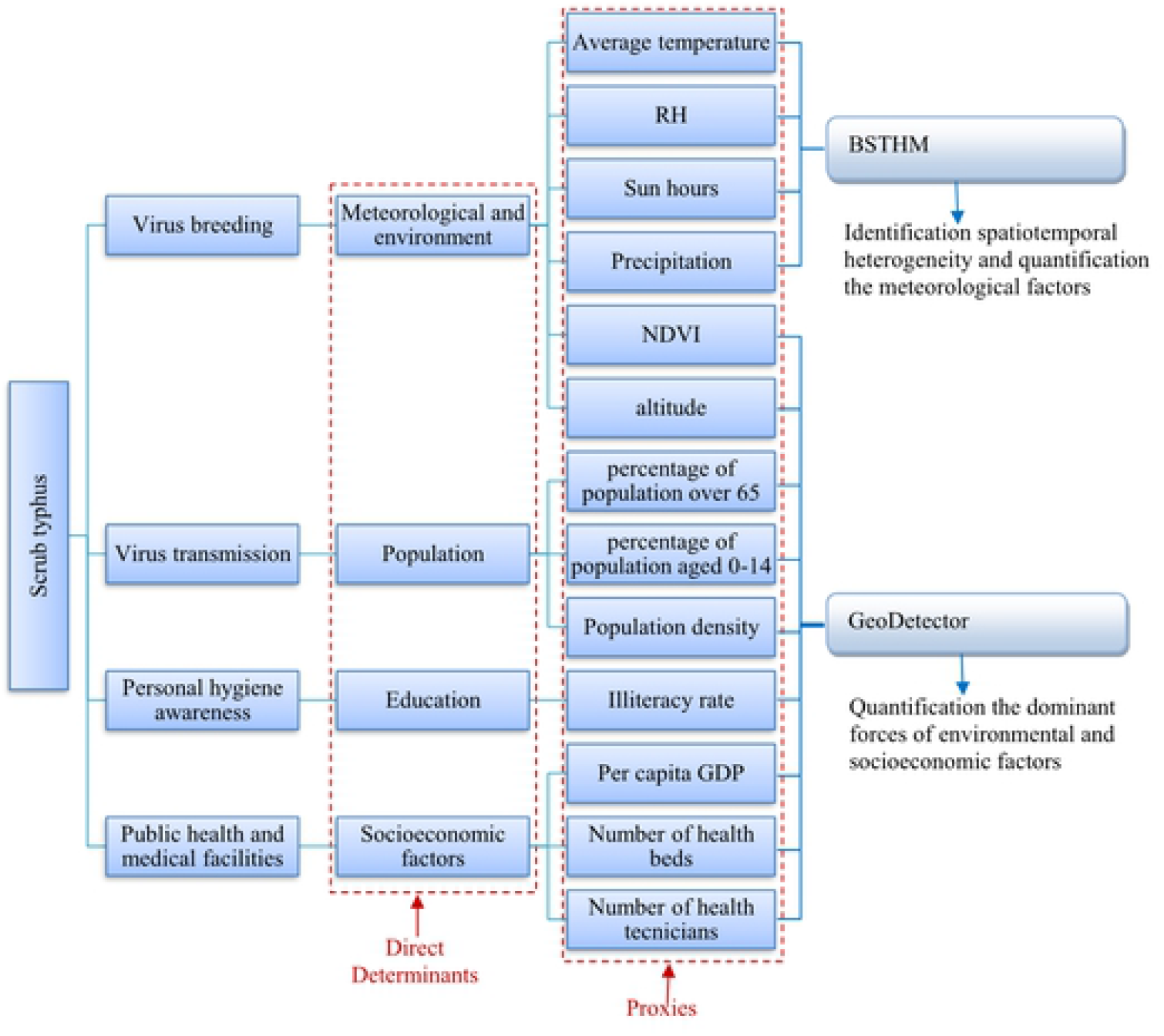
The determinants of scrub typhus and the methods used in this study.

### 2.4 Bayesian space–time hierarchy model (BSTHM)

Space–time phenomena were decomposed into two parts: global and local components. The global part represented common spatiotemporal variation of scrub typhus, whereas the local part revealed the spatiotemporal heterogeneity of the incidence of scrub typhus throughout the whole study period. Specifically, we used the model(Li et al., 2014; Richardson et al., 2004) with the Poisson distribution to capture spatial hotspots and coldspots of scrub typhus and quantity the association between the incidence of this disease and the meteorological factors. In the model, we let *y*_*it*_, *n*_*it*_, and *u*_*it*_ represent the new scrub typhus cases in province *i*(= 1,2,…,31)at time point *t*(= 1,2,…,84), the total population at the end of a year, and the relative risk of scrub typhus incidence, as follows:

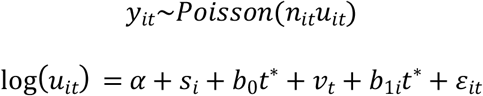

where *α* is the overall log risk of scrub typhus in China and *t*^∗^= *t*―42 (centering at the mid-observation period). The exp(*s*_*i*_)is the spatial risk of this disease, which is influenced by some related factors in the study period, such as economic conditions, local prevention and control policies, and medical resources. (*b*_0_*t*^∗^_+_*v*_*t*_)describes the overall time trend common to all counties with 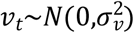. The term *b t*^∗^allows each country to have its own trend. It can be used to capture the deviation from *b*_0_ for each region. For example, if *b*_1*i*_*>*_0_, then the province *i* has a stronger temporal trend than the general trend of the total regions. The last term 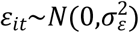 (Gelman, 2006) captures additional variability not yet explained by other model components. A strictly positive half Gaussian prior *N*_+∞_(_0_,1_0_)was given to all random effect standard deviations. In this study, we used the conditional autoregressive (CAR) prior with a space adjacency matrix *W*_31×31_ to impose spatial structure, if the country *i* and *j* shared a common border, then *W*_*ij*_ = 1, otherwise, *W*_*ij*_ = _0_. Based on the posterior parameters of the BSTHM, we classify countries into nine categories(3 risk categories × 3 trend categories) according to a two-stage classification rule(Richardson et al., 2004). In the first stage, a provence is defined as a hotspot if the posterior probaility *p*(*exp*(*s*_*i*_)*>*1|*data*)≥ _0_.8; a provence is defined as a coldspot if *p*(*exp*(*s*_*i*_)*>*1|*data*)*≤*_0_.2; if _0_.2 *< p*(*exp*(*s*_*i*_)*>*1|*data*)*<* _0_.8, the province is defined as neither hotspots nor coldspots. In the second stage, according to the the local slopes *b*_1*i*_, we further classify each risk category in the first stage into three trend patterns: level 1, the increasing in the risk of scrub typhus is faster than the mean trend, if *p*(*b*_1*i*_*>*_0_|*h*_*i*,_*data*)≥ _0_.8; level 2, the increasing in the risk of this disease is slower than the overall trend, if *p*(*b*_1*i*_*>*_0_|*h*_*i*,_*data*)*≤*_0_.8; level 3, the increasing in the disease has no difference with the mean level, if _0_.2 *< p*(*b*_1*i*_*>*_0_|*h*_*i*,_*data*)*<* _0_.8. The whole BSTHM was performed in OpenBUGS (Richardson et al., 2004).

### 2.5 GeoDetector q statistics

GeoDetector is a spatial variance analysis method using the q-statistic, which can be used to quantify the powers of influencing factors and estimate spatio-temporal heterogeneity between scrub typhus and potential risk factors(Wang et al., 2010; Wang et al., 2016; Wang and Xu, 2017). It was expressed as:

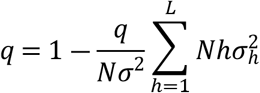

where *q* represents the non-linear relation between the decisive socioeconomic factors and scrub typhus. The value ranges from 0 to 1, and a higher value of *q*-statistic suggesting a higher determinant power of a risk factor or the heterogeneity of a target variable. *N* and *N*_*h*_ are the numbers of provences in the total study area and in the *h*-th stratum(*h* = 1,2,…,*L*),respectively. *σ*^2^ and *σ*^2^ represent the variances in scrub typhus incidence in the entire countries in the *h*-th stratum, respectively.

## 3. Results

### 3.1 Temporal heterogeneity

Over the 84 months between January 2012 and December 2018, a total of 124,341 cases of scrub typhus were reported in the study regions, with a monthly incidence of 0.087 per 100,000 people.. On the time perspective, the overall temporal variation showed an upward trend and a significant seasonality. The incidence of scrub typhus increased significantly in autumn (September to November), with an average monthly incidence of 0.170 per 100,000 people, and reached a peak in October (0.226 per 100,000 people). Meanwhile, the secondary peak appeared in summer (June to August), with an average incidence rate of 0.133/10^5^, and the incidence rate was significantly lower in winter and spring (0.012 per 100,000 people), and the bottom value appeared in February (0.003/10^5^). These results revealed that the risk of scrub typhus had clear temporal stratified heterogeneity, with a *q* statistic value of 0.52.

### 3.2 Spatial heterogeneity

Geographically, the spatial relative risks (RRs) of scrub typhus calculated from BSTHM indicated clear spatial heterogeneity with a *q*-value of 0.64(Fig.3). The provinces with higher spatial RRs mainly appeared in the southern China, while the spatial RRs of northern provinces in China were lower. These results indicated that the southern regions involved in relatively higher risks.

Based on the posterior probability *p*(*exp*(*s*_*i*_)*>*1|*data*), 31 provinces were classified into 3 categories: hot spots, cold spots, and other spots. Among the 31 provinces, 11/31 (35.48%) and 6/31 (19.35%) provinces were identified as cold spots and hot spots, respectively. The remaining 14/31(Wongsantichon et al., 2020 Hainan, Fujian and Jiangxi Hainan, Fujian and Jiangxi), while the cold spots were mainly distributed in the north (Heilongjiang, Jilin, Liaoning, Inner Mongolia, Hebei, Shanxi, Shaanxi, Ningxia, Gansu, Qinghai and Xinjiang)(Fig.4)

Among the six hot spots, the upward trends of Yunnan, Guangdong, Guangxi and Hainan were faster than the overall trend. Consequently, the risk of these regions might be higher than the overall risk and continue to face high incidences in the future. Jiangxi and Fujian showed the same trends as the overall trend, which indicated that these regions would still be hot areas in the future. Therefore, relevant prevention and control departments should focus on these provinces(Fig.4). Among the eleven cold spots, the increasing trend of Xinjiang was the same as the overall trend, so its current risk level would remain constant in the future. The other 10 provinces showed lower increasing than the overall trend, which indicated that the incidence risk of these provinces would be lower than the overall risk and maintain a low-risk state in the future(Fig.4).

Among the remaining 14 provinces of neither hot spots nor cold spots, Beijing, Tianjin, Shanghai, Hubei, and Tibet had a slower upward trend, indicating that these provinces will likely become cold spots in the future, while the other nine provinces were consistent with the overall trend(Fig.4).

The risk of scrub typhus showed obvious seasonal changes and spatial heterogeneity (Fig. 5 and 3), which indicated that meteorological and socio-economic factors played a dominant role in spatiotemporal heterogeneity. In southern China, the factors with the highest determinant powers were average temperature and altitude, while in northern China, the most decisive factors were average temperature and per capita GDP.

**Fig. 3.**
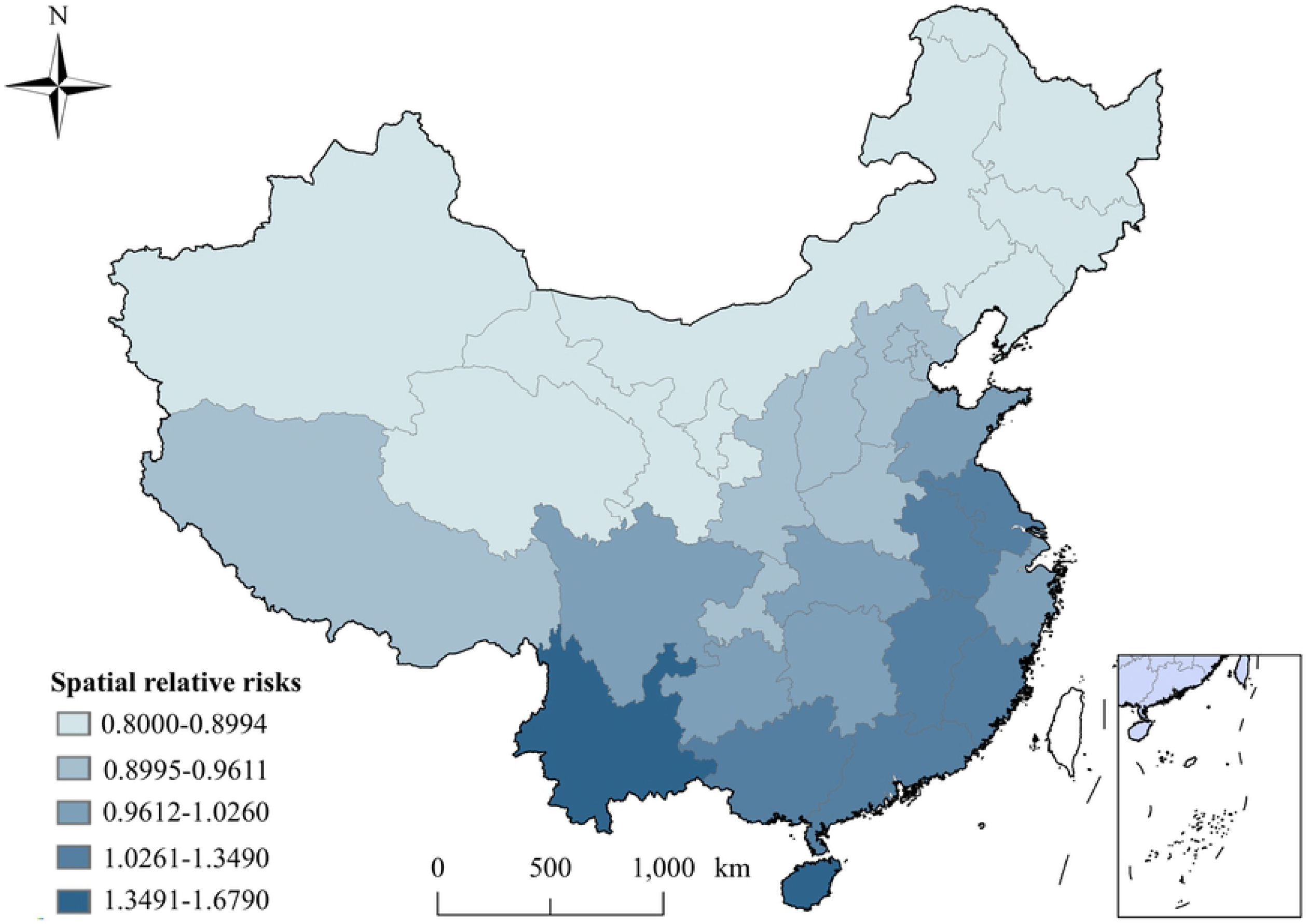
Spatial relative risks of scrub typhus in each province of China.

**Fig. 4.**
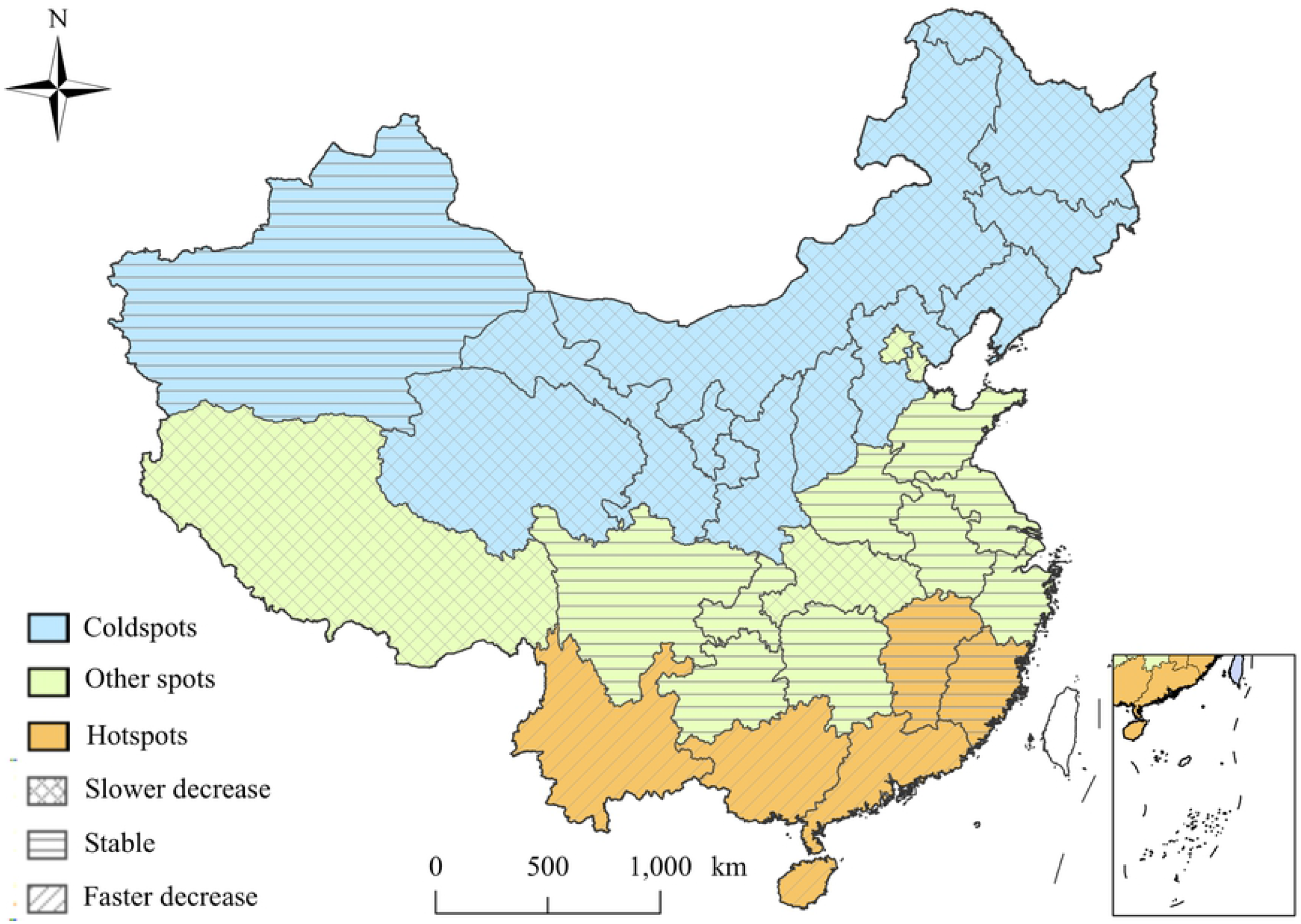
Distribution of hot and cold spots of scrub typhus in China.

**Fig. 5.**
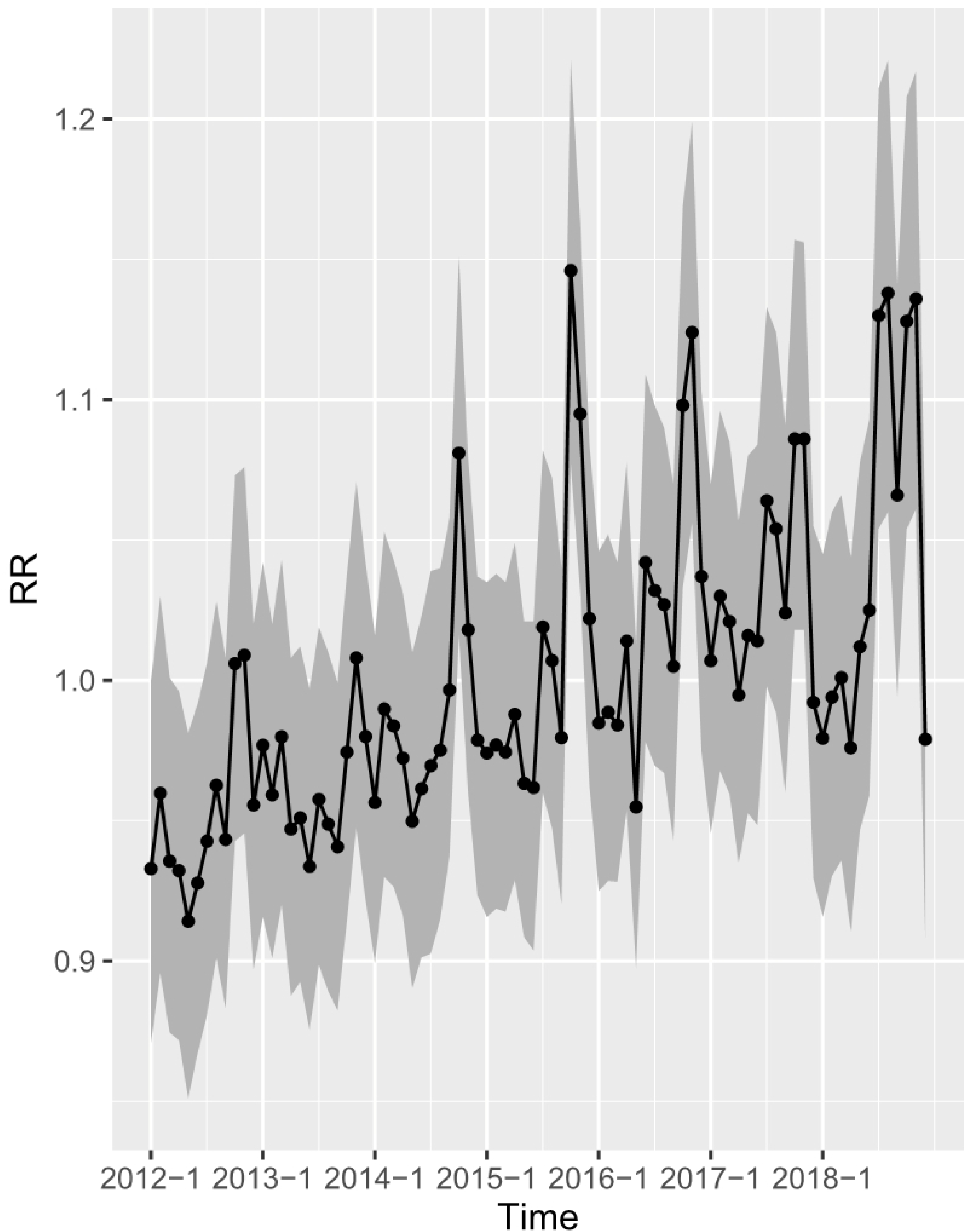
Monthly temporal relative risks of scrub typhus from 2012 to 2018.

### 3.3 Meteorological analysis

Whether in the south or north, the average temperature had a dominant effect on scrub typhus. In the north, the average temperature was positively associated with the risk of scrub typhus. A 1°C increase in the average temperature was associated with an increase of 0.9640% (95% CI: 0.4604-1.4720) in the risk of scrub typhus (RR: 1.0100; 95% CI: 1.0050-1.0150) (Table 2). In contrast, in the south, the average temperature was inversely related to the risk of this disease, and an increase of 1°C in the average temperature was associated with a reduced of 1.1000% (−1.3400, −0.8550) in scrub typhus risk (RR: 0.9891) (Table 1).

The effect of other potential meteorological factors could not be ignored, except for precipitation. For example, in southern China, a 1% increase in RH was associated with a 0.3540% risk reduction (RR: 0.9965). A 1-h increase in total solar hours was related to a 0.0772% increase in scrub typhus risk, with a corresponding RR of 0.9992 (Table 1).

In the north of China, in addition to the average temperature, RH also had a nonnegligible impact on this disease. A 1% increase of RH was related to a 0.7552% increase in scrub typhus risk, with a corresponding RR of 1.0080. A 1 hour increase in total solar hours was associated with an increased of 0.1653% in the risk of scrub typhus (RR: 1.0020). In addition, the estimated coefficients for precipitation was not significant (Table 2).

**Table 1.** The quantified posterior mean values with 95%CI and the relative risks (RRs) for all meteorological factors in the southern provinces of China.

| Meteorological factors | Posterior mean(95%CI) (100%) | RR (95%CI) |
| --- | --- | --- |
| Average temperature(°C) | -1.1000(-1.3400, -0.8550) | 0.9891(0.9867,0.9915) |
| RH (%) | -0.3540(-0.4940, -0.2270) | 0.9965(0.9951,0.9977) |
| Precipitation(mm) | 0.0134(-0.0008, 0.0262) | 1.0000(1.0000,1.0000) |
| Sun hour(h) | -0.0772(-0.1060, -0.0520) | 0.9992(0.9989,0.9995) |

**Table 2.** The quantified posterior mean values with 95%CI and the relative risks (RRs) for all meteorological factors in the northern provinces of China.

| Meteorological factors | Posterior mean(95%CI) (100%) | RR (95%CI) |
| --- | --- | --- |
| Average temperature(°C) | 0.9640(0.4604,1.4720) | 1.0100(1.0050,1.0150) |
| RH (%) | 0.7552(0.5243,0.9438) | 1.0080(1.0050,1.0090) |
| Precipitation(mm) | 0.0287(-0.0201,0.0730) | 1.0000(0.9998,1.0010) |
| Sun hour(h) | 0.1653(0.1114,0.2266) | 1.0020(1.0000,1.0020) |

### 3.4 GeoDetector model

The results of *q* statistics in Geodetector showed that socio-economic and environmental factors also played vital roles in the transmission of scrub typhus. For example, in northern China, per capita GDP was the strongest determinant power, with a *q* value of 0.870. The determinant powers of the number of hospital beds per 1000 people, the number of health technicians, NDVI and altitude were 0.814, 0.789, 0.761, and 0.770, respectively. The determinants of the percentage of population aged 0-14, the illiteracy rate, the percentage of population over 65, and population density were 0.721, 0.684, 0.596, and 0.561, respectively (Table 3).

In southern provinces, altitude was the strongest determinant with a q value of 0.916. The determinants of the percentage of population aged 0-14, illiteracy rate, per capita GDP, and NDVI were 0.882, 0.822, 0.818, and 0.722, respectively. The determinants of the number of health technicians, the number of hospital beds, population density, and the percentage of population over 65 were 0.641, 0.638, 0.233, and 0.221, respectively (Table 3)

**Table 3.** The *q* values (*q* _1_, *q* _2_) calculated for the association between scrub typhus and environmental and socioeconomic factors in northern and southern China, respectively.

| Socioeconomic factors | $q_1$ | $q_2$ |
| --- | --- | --- |
| Normalized Difference Vegetation Index (NDVI) | 0.722** | 0.761** |
| Provincial-level altitude(m) | 0.916** | 0.770** |
| Per capita GDP (10<sup>4</sup>CNY) | 0.818** | 0.870** |
| Population density(person/km<sup>2</sup>) | 0.233** | 0.561** |
| Number of health technicians (per 10<sup>3</sup>) | 0.641** | 0.789** |
| Illiteracy rate (100%) | 0.822** | 0.684** |
| Number of medical beds (bed) | 0.638** | 0.814** |
| Percentage of population aged 0-14(%) | 0.882** | 0.721** |
| Percentage of population over 65(%) | 0.221** | 0.596** |

\*\*Statistical significance level: 0.01.

## 4. Discussion

In recent years, scrub typhus is no longer confined to the Asia Pacific region and has become a public health problem worldwide (Costa et al., 2021; Silva-Ramos et al., 2021; Weitzel et al., 2019). In China, the incidence of scrub typhus increased steadily before 2012 and increased rapidly but unevenly after 2012(Sun et al., 2017; Yu et al., 2018). In this study, BSTHM was used to analyze the spatiotemporal heterogeneity of scrub typhus risk and to quantify the impact of meteorological factors on scrub typhus from 2012 to 2018. The influence of socioeconomic and environmental factors on scrub typhus was estimated using GeoDetector model in southern and northern China. The results showed that the spatial distribution of scrub typhus in China was inhomogeneous. The highest risk provinces (hot spots) of this disease were mainly distributed in the south, while the lowest risk areas were clearly distributed in northern China, thus indicating that the difference of environmental and socio-economic factors were closely associated with scrub typhus. The average temperature and per capita GDP were the determinant factors in the north, while average temperature and altitude were the dominant influencing factors in the southern China.

The spatial heterogeneity may be attributed to, but not limited to, the following reasons. On the one hand, regional differences may lead to differences in vectors and hosts between the two regions, such as the differences of the number and species of Chigger Mites and rodents. The complex and diverse terrain and high biodiversity of the landscape in the South provide a suitable habitat for Chigger Mites and rodent hosts(Derne et al., 2015). For example, in China, the hosts of Orientia tsutsugamushi are mainly Rattus losea, R. tanezumi, R. norvegicus, Nivirenter confucianus and Apodemus agrarius(GuangHua et al., 2013), of which Rattus losea and R. tanezumi are mainly distributed in the south. Chigger Mites are both hosts and vectors, mostly distributed in provinces and regions from the southeast coast to the southwest border of China.

On the other hand, the influence of socio-economic factors are nonnegligible. With the development of tourism and the construction of urban and rural areas, the living environment of rodents has changed, resulting in a large number of migration of rodents, which may be one of the reasons for the emergence of new scrub typhus foci. Some studies had shown that the expansion of urban parks and greening has increased the risk of residents suffering from scrub typhus(Park et al., 2015; Wei et al., 2014). Therefore, strengthening the monitoring of host animals, Chigger Mites and Orientia tsutsugamushi infection is conducive to preventing and controlling the spread of Orientia tsutsugamushi disease and reducing the incidence.

Our study showed that the percentage of children younger than 14 years old and the illiteracy rate were important factors affecting the risk of scrub typhus. A study on the Penghu Islands also showed that higher school enrolment seemed to be associated with a reduction of child disease infection rate(Olson and Bourgeois, 1979). Meanwhile, scattered children and students have always been in the top 4 on the occupational list of scrub typhus, which means that children are a neglected group, and we need to strengthen their education and pay attention to preventive measures(Yao et al., 2019). The risk of scrub typhus also presented obvious temporal heterogeneity.The high-risk seasons were summer (June to August) and autumn (September to November), especially autumn, which was consistent with the results observed in some previous studies(Ren et al., 2019; Yang et al., 2021). It may be related to the dominant vector and animal host of scrub typhus. For example, the reproductive peak of Rattus losea is from March to October, with the highest in October, and the lowest is in February, which is similar to the case change of scrub typhus. Leptotrombidium deliense is the main vector of summer type scrub typhus, and leptotrombidium scutellare is the main vector of autumn winter type scrub typhus(Wu et al., 2013). This seasonal change will help the public health department to formulate reasonable prevention and control policies and allocate resources according to the peak incidence and the months with more reported cases reasonably.

Meteorological factors are considered to be important environmental factors.This study showed that temperature, RH and sunshine hours were factors playing predominant roles affecting the transmission of scrub typhus, which is consistent with some previous studies(Yang et al., 2014; Zheng et al., 2019). In the north, the average temperature, RH and sunshine hours were positively correlated with scrub typhus risks. On the contrary, in the south, the average temperature, RH and sunshine hours showed a negative association with scrub typhus, indicating that the meteorological range is a decisive factor.

Our study identified average temperature as positively associated in northern China and negatively correlation in southern China with scrub typhus risk, revealing that the temperature was a significant factor on scrub typhus incidences, which pandered to the previous studies on infectious diseases, such as hand-foot-and-mouth disease(Hu et al., 2020), influenza(Park et al., 2020), and scrub typhus(Lu et al., 2021). Temperature may affect the spawning rate and activity of chiggeridae and the abundance of hosts. Lu et al.’ research showed that before 29.6 °C, with the increase of temperature, the oviposition rate of chiggeridae and the number of rodents increased, and Chigger Mites became more active(Van Peenen et al., 1976). The RR value was the highest at 27°C, and the weekly temperature range was negatively correlated with the risk of disease(Lu et al., 2021). When the temperature is higher, people will reduce their willingness to farm or go out (such as parks), thus reducing the risk of tsutsugamushi infection.

Monthly average RH was negatively correlated with the risk of scrub typhus in the south, which is different from expection, because higher RH means more mites. This negative correlation can be attributed, but not limited, to the following two reasons: One is that the water source for mites to survive is water vapor(Clopton and Gold, 1993), so RH has always been a crucial factor in determining the number of Chigger Mites. However, higher humidity is also not conducive to the life cycle of Chigger mite. For example, Yao et al. ‘study(Yao et al., 2019) found that areas with relative humidity higher than 63% have lower risk than other areas. Therefore, a detailed monitoring of environmental humidity possibly helped to provide a fine map of future risk of scrub typhus. Secondly, the negative correlation might be related to the interaction between human and environment. For example, in the south, people always prefer to engage in outdoor activities in the dry season, thus giving more exposure opportunities to Chigger Mites(Zheng et al., 2019). Therefore, when carrying out outdoor activities, we should improve our awareness of prevention and take appropriate protective measures, such as don’t lie directly on the grass, but lay blankets or wear protective clothing.

Some previous studies showed that sunshine was also a dominant factors determining the activity time of Chigger mite larvae(Clopton and Gold, 1993), which may partly explain the lower incidence in northern China than in the south. The results showed that the number of sunshine hours was positively correlated with scrub typhus within a certain range, but showed inhibitory effect when it exceeded this range. For example, in Yao et al. ‘(Yao et al., 2019) BRT model, sunshine hours of less than 170 was positively correlated with the risk of scrub typhus, while sunshine hours more than 170 was negatively correlated with the disease.

This study has some limitations. First of all, we used provincial data to explore the association at the group level, which may lead to an ecological fallacy inevitablely, but this did not influence the long-term trends of scrub typhus. Secondly, as a multifactorial disease, factors other than those considered in this study may have introduced some uncertainties on scrub typhus. And then, new factors such as the prevalence of emerging infectious diseases will also affect the spread of scrub typhus.

## Competing interest

The authors declare that they have no known competing financial interests or personal relationships that could have appeared to influence the work reported in this paper.

## Author Contributions

**Funding acquisition:** Changqiang Zhu, Weilong Tan.

**Data accept:** Jiaojiao Qian, Lele Ai, Fuqiang Ye, Yong Qi.

**Data analysis:** Jiaojiao Qian, Hongliang Chu, Ji He, Chongcai Wang.

**Project administration:** Jing Yi, Chunhui Wang, Weilong Long.

**Supervision:** Chunhui Wang, Weilong Long.

**Writing-original draft:** Jiaojiao Qian, Weilong Tan.

**Writing-review&editing:** Jing Yi, Jiaojiao Qian, Weilong Tan.

## Notes

### Competing Interest Statement

The authors have declared no competing interest.

